# Inflammation-profiling reveals activated pathways and biomarkers with predictive potential in oligoarticular JIA

**DOI:** 10.1101/2025.03.17.643682

**Authors:** Xingzhao Wen, Cecilia Aulin, Erik Sundberg, Heshuang Qu, André Struglics, Anne-Sophie Merritt, Erik Melén, Maria Altman, Helena Erlandsson Harris

**Author notes:** Correspondence to: Prof. Helena Erlandsson Harris, Center for Molecular Medicine, division of Rheumatology, Department of Medicine Solna, Karolinska Institutet and Karolinska University Hospital, SE-171 76 Stockholm.

## Abstract

**Objective:** We set out to profile the immune mechanisms active in treatment-naïve oligoarticular JIA (oJIA) to improve understanding of its immunopathogenesis, to identify potential biomarkers that can aid diagnosis, predictions and correlate with clinical disease parameters.

**Methods:** Using Olink proteomics (Inflammation Panel), we defined and compared the inflammation profiles of 38 plasma and 62 synovial fluid (SF) oJIA samples, 38 plasma samples from healthy age- and sex-matched controls (HC), 12 SF samples from non-arthritic controls and 26 SF samples from knee injury patients. Clinical data were retrieved from the Swedish Pediatric Rheumatology Quality Register.

**Results:** Plasma profiles of oJIA and HC were largely overlapping, with IL6 and MMP-1 upregulated in oJIA. In SF, 48 differentially expressed proteins (DEPs) were identified in oJIA, highlighting immune pathways like leukocyte migration, cell chemotaxis and adaptive immunity. Comparative analysis revealed 13 proteins specific to oJIA. Correlations were found between DEPs in oJIA SF and clinical parameters (cJADAS-71, pain, health impact score). Plasma IL6 and MMP-1 showed strong correlation with disease activity and pain, respectively. CXCL9, CXCL10 and CXCL11 were identified as potential predictive biomarkers for disease progression.

**Conclusions:** The overlap in plasma inflammation profiles of oJIA and HCs suggests local rather than systemic inflammation in oJIA, underlining the need for synovial fluid-based immunopathogenesis studies. Adaptive immune signatures in oJIA SF distinguished it from knee injury patients, offering potential for diagnostic application. Increased CXCL9, CXCL10 and CXCL11 in SF were associated with chronic disease progression and could serve as prognostic biomarkers and early treatment targets.

## 1. Introduction

Juvenile idiopathic arthritis (JIA) is a heterogeneous group of chronic childhood arthritis of unknown etiology, characterized by arthritis persisting for more than six weeks with onset before the age of 16 [1]. The incidence of JIA ranges from 1.6 to 23 cases per 100,000 children [2, 3]. The International League Against Rheumatism (ILAR) has identified seven subtypes of JIA according to disease manifestations within the initial 6 months of the disease, with oligoarticular arthritis representing the largest subtype affecting half of all JIA cases [1, 4].

Since the introduction of biological therapies three decades ago, treatment success has improved substantially for children with JIA but a cure is still not available. Precise diagnosis, assessment, and treatment for JIA patients in the early stage are critical to the outcomes [5]. Thus, it is important for clinical doctors to stratify patients in advance for subsequent targeted follow-up and treatment decision-making. These decisions are hampered by the still existing knowledge gaps of the complex immunopathological mechanisms driving the different subtypes of JIA and the different hallmarks of disease; inflammation, pain, fatigue and joint destruction[6].

Recently, several biomarkers that could improve the diagnosis of JIA subtypes and prediction of disease trajectories have been identified. For instance, S100A8/A9 and S100A12 can serve as diagnostic biomarkers and are associated with response to anti-tumor necrosis factor (TNF) treatment in systemic JIA patients [7, 8].14-3-3η has been found to have the potential to serve as a new biomarker for polyarticular JIA [9]. Most of the suggested biomarkers reflect the inflammatory disease status, only a few have been suggested to reflect the other hallmarks of the disease. Surprisingly, only few biomarkers have been proposed for oJIA, despite it being the most common JIA subtype. Raggi et al. reported microRNAs and proteins derived from extracellular vesicle have the potential to serve as early molecular indicators of oJIA [10, 11]. Despite the invested efforts, there is still a dire need to identify new and precise diagnostic and prognostic biomarkers as well as measurable biomarkers that reflect clinical features.

Proteins related to inflammation hold significant potential to serve as diagnostic and prognostic biomarkers in oJIA [12]. Studies reporting comprehensive inflammation profiles of oJIA patients are few and the use of local tissue samples are rare. Moreover, treatments such as DMARDs and biological DMARDs could dramatically alter the inflammation profiles [13]. Therefore, there is an urgent need to obtain early and treatment naïve inflammation profiles of oJIA patients.

In this study, we used plasma and synovial fluid (SF) samples from treatment naïve oJIA patients with short disease duration for cross-sectional proteomics analysis and described the inflammation profiles. We described how they differ from healthy controls and from subjects with traumatic knee injuries. We identified proteins that could serve as biomarkers for diagnostic and prognostic purposes, that correlate with clinical hallmarks of disease and should be investigated as targets for therapy development.

## 2. Methods and Materials

### 2.1 Study population and data sources

Human subjects research was approved by the North Ethical Committee in Stockholm, Sweden (Dnrs 2009–1139-01 and 2010–165–31 for Juvenile Arthritis BioBank Astrid Lindgren’s hospital and Dnr 03–067 for Barnens miljö-och hälsoundersökning). Plasma (n=38) and SF (n=62) samples from 63 different oJIA patients (37 paired plasma SF samples) were collected at Astrid Lindgren’s Children Hospital, Karolinska University Hospital, Stockholm, Sweden. These patients all met the ILAR criteria and were either on no medication or treated with NSAIDs only at the time of sampling. The disease duration (from onset to sampling) was less than two years. Clinical and laboratory data collected at each sampling occasion were retrieved from medical charts and from the Swedish Pediatric Rheumatology Quality Register (Svenska Barnreumaregistret, Omda®) and presented in table 1. Plasma samples from 38 age- and sex-matched healthy individuals were selected from a population-based cohort (Barnens miljö-och hälsoundersökning [14]) from the Stockholm region. Juvenile control SF samples from 12 individuals who underwent arthroscopic examination three months after a knee injury, during which no abnormal clinical features were observed, were collected in the Lund region, Sweden [15]. From the same Lund region cohort, SF samples from 26 juvenile patients with acute knee injuries were collected during arthroscopic examination and selected for analysis. The main clinical information of the study cohorts is presented in Table 1. We also used a data set from a publicly available, independent oJIA cohort [16] of SF samples from 41 oJIA patients to validate the prognostic biomarkers identified in our analysis (Table S1).

**Table 1.** Demographics and disease characteristics of the study population.

|  | oJIA patients<br>(plasma) | Healthy individuals<br>(plasma) | oJIA patients<br>(SF) | Juvenile control<br>(SF) | Juvenile knee injury<br>(SF) |
| --- | --- | --- | --- | --- | --- |
| Sample size(n) | 38 | 38 | 62 | 12 | 26 |
| Sex(F) | 22 | 22 | 38 | 8 | 5 |
|  |  | *P*=1 (vs oJIA plasma) |  | *P*=0.979 (vs oJIA SF) | *P*<0.001 (vs oJIA SF) |
| Age at sampling (median/range) | 7.5 (1-15) | 8 (4-12) | 9 (1-16) | 14 (11-16) | 16 (13-18) |
|  |  | *P*=0.467 (vs oJIA plasma) |  | *P*<0.001 (vs oJIA SF) | *P*<0.001 (vs oJIA SF) |
| **CRP** Measurement(n) | 29 |  | 47 |  |  |
| median/range (mg/L) | 3 (1-38) |  | 3 (<1-60) |  |  |
| **ESR** Measurement(n) | 29 |  | 42 |  |  |
| median/range (mm/h) | 12 (3-97) |  | 11 (2-97) |  |  |
| **RF** Measurement(n) | 18 |  | 38 |  |  |
| positive | 1 (5.56%) |  | 1 (2.63%) |  |  |
| **ANA** Measurement(n) | 35 |  | 61 |  |  |
| positive | 17 (48.57%) |  | 27 (44.26%) |  |  |
| **HLA-B27** Measurement(n) | 5 |  | 16 |  |  |
| positive | 1 (20%) |  | 3 (18.75%) |  |  |
| Active joint count (median/range) | 1 (1-5)* |  | 1 (1-5) # |  |  |
| cJADAS-71 | 6.07±3.09* |  | 5.86±3.43# |  |  |
| VAS pain | 3.97±2.45* |  | 3.68±2.22# |  |  |
| Health impact(median/range) | 2.20 (0.10-9.00) * |  | 2.70(0.10-9.00) # |  |  |

\*: 5 patients lack the scores of these clinical parameters, resulting in 33 patients measured in total.
#: 7 patients lack the scores of these clinical parameters, resulting in 55 patients measured in total.
Abbreviation: C-reactive protein (CRP); Erythrocyte sedimentation rate (ESR); Rheumatoid factor (RF); Antinuclear antibodies (ANA); Human leukocyte antigen B27 (HLA-B27); Clinical Juvenile Arthritis Disease Activity Score of 71 points (cJADAS-71); patient visual analogue scale of pain (VAS pain,0-10); patient visual analogue scale of well-being (health impact,0-10)

### 2.2 Sample Collection and Processing

Fresh SF and blood samples were collected in citrate tubes and ethylenediaminetetraacetic acid (EDTA) tubes separately. SF and blood samples were centrifuged at 3000g for 10-15 min to obtain cell-free samples. All samples were prepared and aliquoted within 4 h and stored at −80L until analysis. SF samples were diluted with PBS in a 1:4 ratio for Olink proteomics analysis based on our previous study [15].

### 2.3 Proteomic assay

Plasma and SF samples were sent for analysis by a high-throughput multiplex immunoassay (Target 96 Inflammation Panel, Olink Bioscience, Sweden). The immunoassay is based on the proximity extension assay technology [17] and the inflammation panel includes 92 proteins associated with inflammatory and immune response processes (https://olink.com/products/olink-target-96). The protein levels are recorded as log2-normalized protein expression (NPX) values.

### 2.4 Quality control and data pre-processing

Data for proteins with NPX values below the limit of detection (LOD) in more than 80% of the samples were removed. Consequently, 84 proteins in the plasma samples and 76 proteins in the SF samples were included in further analysis.

The plasma samples included in this study were run in two versions of the Olink Target 96 inflammation panel and the batch correction method ComBat was applied to normalize the log base 2 normalized protein expression (NPX) values between the two runs. Data for plasma IFN-gamma and TNF levels were excluded from the analysis as the antibodies included in the panel had been changed between the two runs. Finally, values for 82 proteins in plasma and 76 proteins in SF were included in the ensuing analyses. Additional details of the 92 proteins in the Olink panel, including the abbreviated names are listed in Table S2.

### 2.5 Data processing

Principal Component Analysis (PCA) and Hierarchical Clustering Analysis (HCA) were performed by ClustVis Web Tool (https://biit.cs.ut.ee/clustvis/).

Differentially expressed proteins (DEPs) between groups were defined as ΔNPX higher than 1 (fold change >2) and a p-value of less than 0.05. The p-values were corrected with Benjamini-Hochberg method for multiple comparisons to control the false discovery rate (FDR) at 5% within each comparison.

STRING database 12.0 was used to construct and visualize the protein-protein interaction network and enriched pathways for DEPs. Additionally, gene ontology (GO) biological process pathway analysis of DEPs was performed to identify enriched pathways.

### 2.6 Statistical analysis

IBM SPSS Statistics 25 software and R (R 4.3.2) were used for statistical analyses. Distribution of protein expression and clinical assessment data were assessed using Shapiro-Wilk test (n<50) or Kolmogorov-Smirnov test (n>50). Correlation analyses were performed using Pearson or Spearman methods, depending on data distribution. A p-value less than 0.05 was considered statistically significant. Random forest classification and factors’ importance calculations were performed using the random forest package in R. Receiver operating characteristic curves (ROC) were analyzed with the pROC package.

## 3. Results

### 3.1 Overlapping Plasma inflammation profiles in oJIA patients and healthy controls

We first defined and compared the plasma inflammation profiles of oJIA patients and age- and sex-matched healthy controls (HC). PCA and non-supervised HCA revealed a major overlap of the inflammation profiles (Figure 1A-1B), suggesting similar protein expression patterns in both groups. Neither sex nor age resulted in a separation of the two groups (Figure 1B). IL6 and MMP-1 were significantly higher in oJIA group compared to HC (Figure 1C-1D). Unexpectedly, 10 inflammation-related proteins, SIRT2, ST1A1, CXCL5, CXCL6, AXIN1, CASP-8, CXCL1, STAMBP, 4E-BP1 and IL7, showed lower expression levels in oJIA patients than in HC (Figure 1C-1D, Table S3). The number of DEPs identified in plasma between oJIA and HC constrains the ability to study the pathogenesis of oJIA using the plasma proteome.

**Figure 1.**
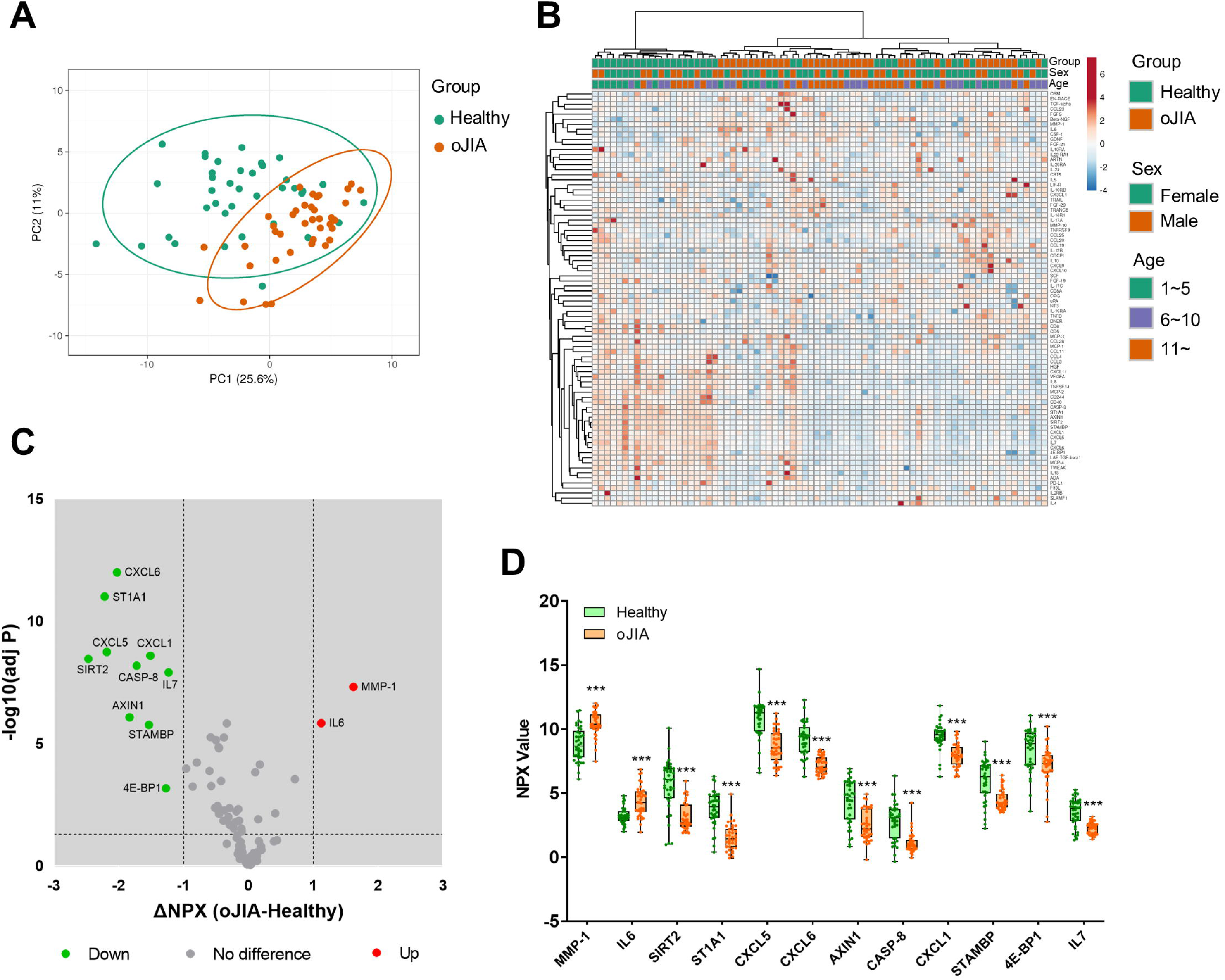
Plasma inflammation profiles in oJIA (n=38) vs HC (n=38) groups. A) PCA results of the oJIA and HC groups based on 82 included proteins in plasma. Confidence level of the ellipses is 0.95. Each point represents a single patient, with a total number is 76 including 38 oJIA and 38 HC. B) Heatmap visualization of 82 included proteins expression in oJIA and HC group, with unsupervised hierarchical clustering analysis revealing an overlap between these two groups. C) Volcano plot shows DEPs in oJIA compared to HC. Broken lines indicate significance thresholds (adjusted PL<L0.05 and ΔNPX >1). D) Boxplot presents 12 DEPs between oJIA and HC group. Statistics: D unpaired t test with correction of multiple comparison by Benjamini-Hochberg method, controlling the FDR at 5%. * P < 0.05, ** P < 0.01, *** P < 0.001

### 3.2 Discrete synovial fluid inflammation profiles in oJIA patients and controls

To further reveal pathogenic mechanisms in oJIA at the site of the main target tissue, the joint, the inflammation profiles of SF from oJIA patients and controls were determined. Given that the mean age of the control group was significantly higher than that of the oJIA group, posing a potential confounding factor, the analysis was initially restricted to oJIA patients aged 11 or older to ensure a better age matching. This initial analysis yielded results consistent with results obtained when including all oJIA patients and controls (Figure S1A-1B and Figure 2A-2C). PCA conducted to assess the inflammation profiles across different age groups, revealed a substantial overlap (Figure S1C). Considering the negligible impact of age on the observed profiles, data from all oJIA patients and controls were included in subsequent analyses to enhance the statistical power.

**Figure 2.**
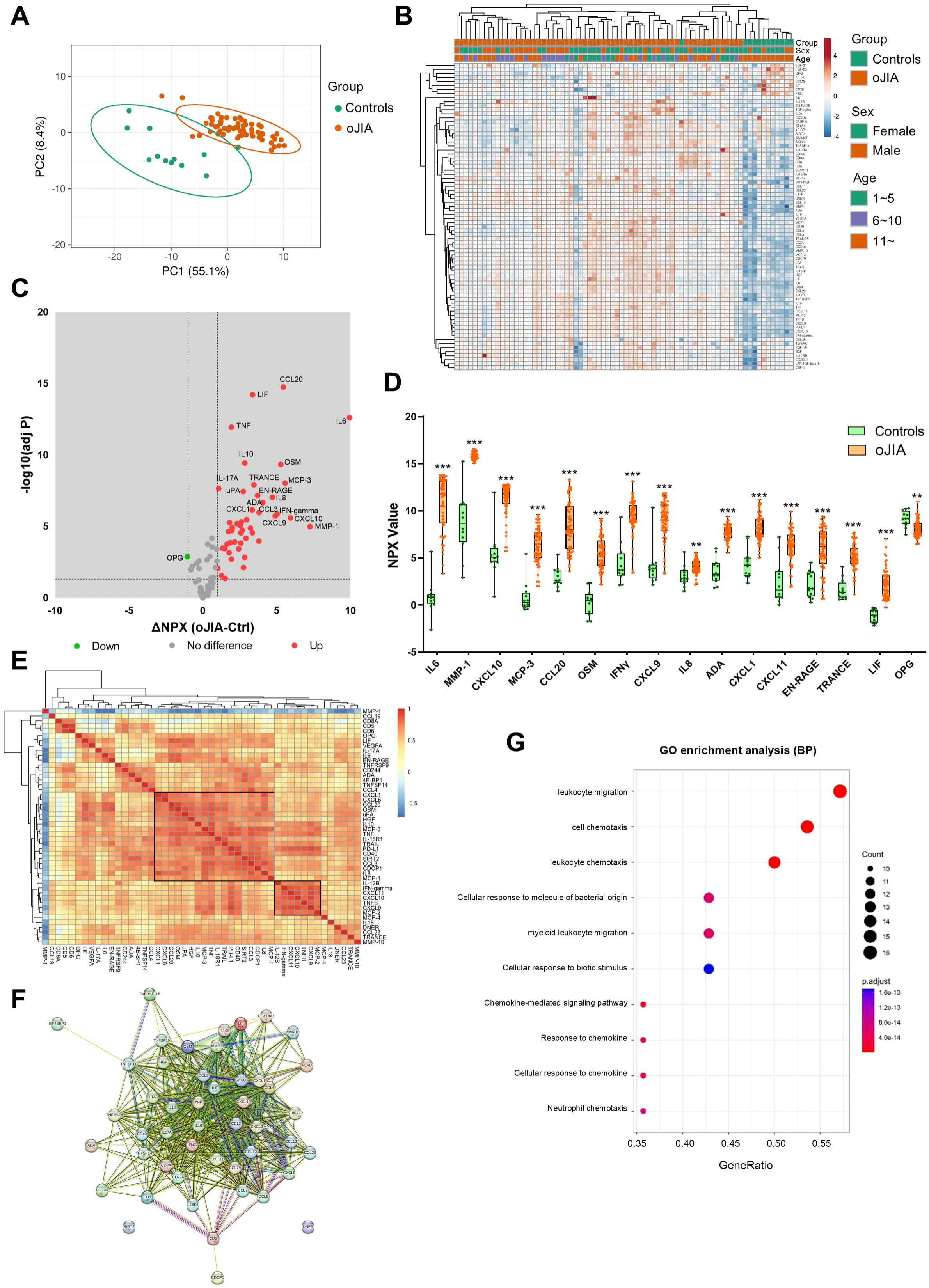
SF inflammation profiles in oJIA patients (n=62) vs controls (n=12) groups. A) PCA results of the oJIA and control groups based on 76 included proteins in SF present a separation between two groups. Confidence level of the ellipses is 0.95. Each point represents a single patient, with a total number is 74 including 62 oJIA and 12 controls. B) Heatmap visualization of 76 included proteins expression in oJIA and control group, and unsupervised hierarchical clustering analysis reveal a separation between oJIA and control. C) Volcano plot show DEPs in oJIA compared to controls. Broken lines indicate significance thresholds. D) Top 15 increased proteins in OJIA and OPG, which is only one decreased protein in oJIA are presented with boxplot. E) Heatmap visualization of the results of correlations (Pearson r) of DEPs in oJIA patients (n=62). The right label illustrates the scores of Pearson r: The closer r is to 1, the redder the color becomes. F) The results of STRING analysis based on DEPs in oJIA reveal hub proteins and enriched pathway. G) GO enrichment analysis based on the DEPs in oJIA highlights leukocyte migration, cell chemotaxis, chemokine mediate signaling, etc involved in pathogenesis of oJIA. Statistics: D unpaired t test with correction of multiple comparison by Benjamini-Hochberg method, controlling the FDR at 5%. E Pearson correlation. * P < 0.05, ** P < 0.01, *** P < 0.001

PCA and non-supervised HCA revealed a discrete separation of the SF inflammation profiles in oJIA patients and controls (Figure 2A-2B). 47 proteins including IL6, MMP-1 and CXCL10 were significantly increased in oJIA and only one protein, OPG, was decreased in oJIA (Figure 2C, Table S4). The top 15 increased proteins as ranked by ΔNPX and OPG are presented in Figure 2D. To further investigate the interactions of the SF DEPs, we performed correlation analysis to investigate the relationship of expression of each DEP in oligo patients and STRING analysis to identify the interacted network. Correlation heatmap showed a few clusters of proteins that correlated with each other (Figure 2E), such as a cluster of 18 proteins including CXCL1, CXCL6, CCL20 and OSM, and a cluster of 7 proteins including IL-12B, IFN-gamma, CXCL11, CXCL10, TNFB, and CXCL9. STRING network showed IL6, TNF, IL10, and IFN-gamma were hub proteins in the interactions network of DEPs and most DEPs are involved in chemokine interactions (Figure 2F). GO enrichment analysis also revealed these DEPs play roles in leukocyte migration, cell chemotaxis and chemokine mediate signaling (Figure 2G). Taken together, oJIA patients exhibit a distinct immunoprofile in SF compared to controls, characterized by an increase in inflammatory proteins. In addition, leukocyte migration and chemokine interactions play important roles in the pathogenesis of oJIA.

### 3.3 A different inflammation pattern in oJIA compared to acute inflammation

We further compared the inflammation profiles of oJIA, knee injury, and controls in SF samples. Each group displayed a significant separation from the other two groups displayed in the PCA plot (Figure 3A), similar to the three separate clusters defined by non-supervised HCA (Figure 3B). Volcano plot showed 45 proteins were significantly increased in knee injury SF compared to controls (Figure 3C). We then compared the increased proteins in oJIA with the increased proteins in knee injury, and found 13 oJIA-specific proteins, 11 injury-specific proteins and 34 overlapping proteins. For example, IL6, MMP-1, CCL20 and OSM commonly increased in both oJIA and knee injury, while ADA, IFN-gamma and TNF specifically increased in oJIA, and CASP-8, CCL11 and SCF specifically increased in knee injury patients (Figure 3D). To further investigate the differences between oJIA and knee injury group, we applied STRING analysis on the unique increased proteins in each group. Adaptive immunity, TNF binding interactions, and T cell modulation were implicated in the pathogenesis of oJIA, while knee injury was characterized by activation of the PI3K pathway (Figure 3E-3F).

**Figure 3.**
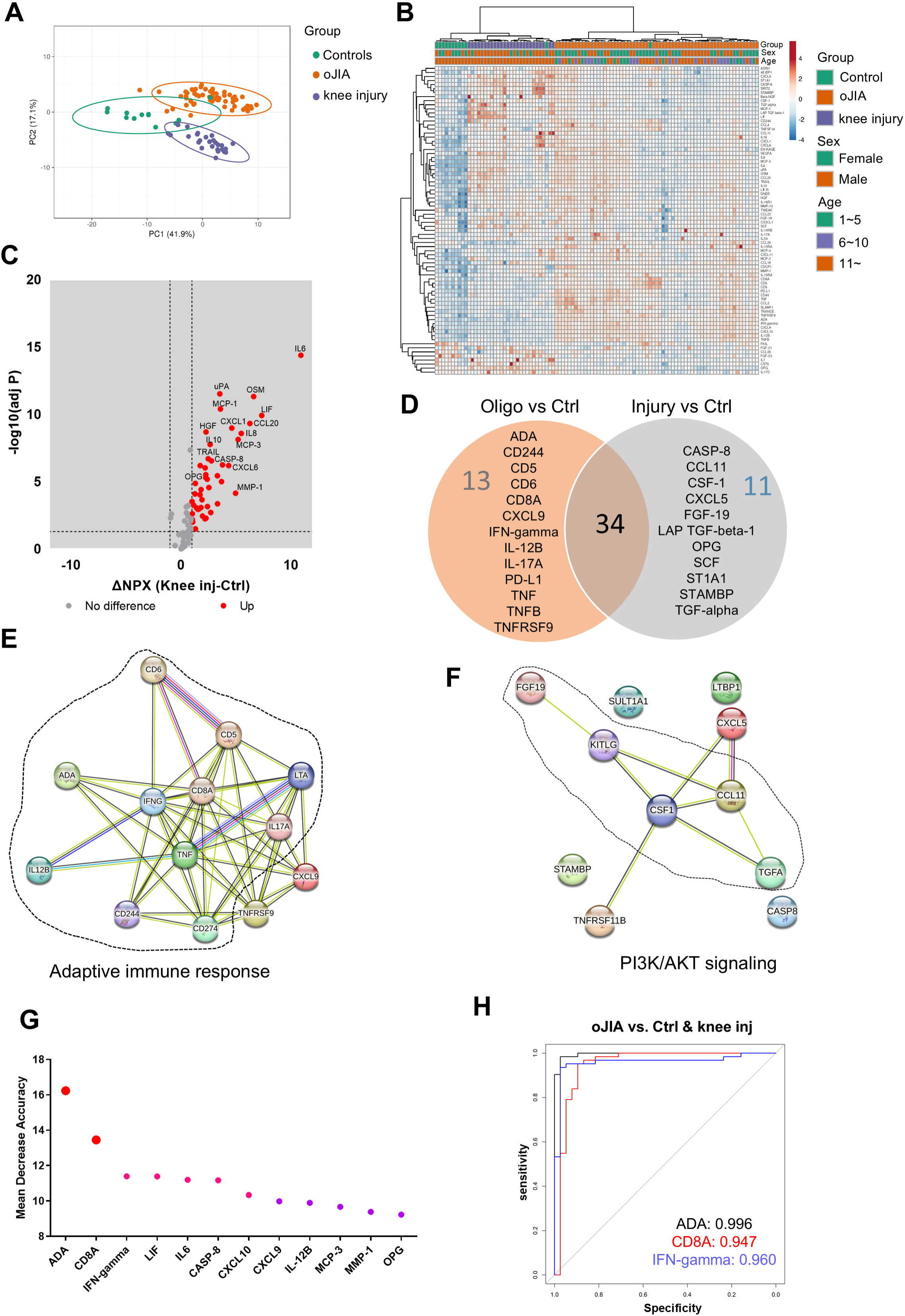
oJIA patients have a different inflammation pattern from knee injury. A) PCA results of the oJIA, knee injury, and control groups based on 76 included proteins in SF present a separation among these three groups. Confidence level of the ellipses is 0.95. Each point represents a single patient, with a total number is 100 including 62 oJIA, 26 knee injury and 12 controls. B) Heatmap visualization of 76 included proteins expression in oJIA, knee injury, and control group, and unsupervised hierarchical clustering analysis reveal a separation among three groups. C) Volcano plot show DEPs in knee injury compared to controls. Broken lines indicate significance thresholds. D) Venn plot comparing the increased proteins in oJIA and those in knee injury compared to controls reveal 34 commonly increased proteins, 13 specifically increased proteins in oJIA, and 11 proteins in knee injury. E-F) The results of STRING analysis based on specifically increased proteins in oJIA or in knee injury reveal enriched pathway. G) Random Forest analysis among three groups highlights proteins contributing the separation. H) ROC analysis based on ADA, CD8A, and IFN-gamma shows these three proteins have great potential to distinguish oJIA from injury and healthy individuals.

To identify the important proteins that contributed to the separation between oJIA, knee injury, and controls, we performed random forest (RF) classification. Here we could discern ADA, CD8A, and IFN-gamma ranked top three in the important proteins contributing to the separation (Figure 3G). To further investigate their individual capability of distinguishing oJIA patients from knee injury and control group, ROC analysis was performed. The results showed that ADA (AUC:0.996, P<0.001, 95%CI: 0.990-1), CD8A (AUC:0.947, P<0.001, 95%CI: 0.890-1), and IFN-gamma (AUC:0.960, P<0.001, 95%CI: 0.917-1) were best in distinguishing oJIA groups from juvenile knee injury and control group, while the other markers (from Figure 3G) had AUC between 0.565 – 0.868 (Figure 3H, Table S5).

### 3.4 Correlation with DEPs in SF vs plasma

To better understand the association between local markers of inflammation and how/if these are reflected systemically, we performed correlation analysis between the DEPs in SF between oJIA and controls and the same proteins in plasma. Seven proteins showed significant correlation (|r|≥0.4, P<0.05) between SF and plasma: IL6, MMP-10, EN-RAGE (S100A12), MMP-1, VEGFA, CCL4, and 4E-BP1 (Table S6). The scatter plots presented the distribution and correlation of these proteinsin SF and plasma (Figure S2A-S2G). ADA and CD8A did not show significant correlation between SF and plasma, indicating that they could work as markers in SF but not as systemic (Figure S2H-S2I).

### 3.5 Assessments of DEPs and oJIA clinical parameters

To evaluate the clinical application potential of the DEPs identified in our study, we investigated the correlations between DEPs and clinical assessment parameters in both SF and plasma. The clinical parameters included cJADAS-71, pain, active joint number and patient reported pain and health impact. The data was available from 55 of the oJIA patients, corresponding to 54 SF and 33 plasma samples.

In SF, 22 proteins correlated with cJADAS-71 score (|r|≥0.4, P<0.05), where CCL20, IL6 and OSM displayed strong correlation (r>0.7) (Figure 4A, Table S7). Interestingly, MMP-1 was increased in oJIA but had a negative correlation with cJADAS-71 (Figure 4A). 12 proteins including IL6, MMP-1, CCL20 show moderate correlation with pain (Figure 4A, Table S7). And 6 proteins correlate moderately with health impact (Figure 4A, Table S7). However, none of proteins in SF correlated with the active joint number. Full datasets of the correlation between DEPs in SF and clinical parameters are listed in Table S7.

**Figure 4.**
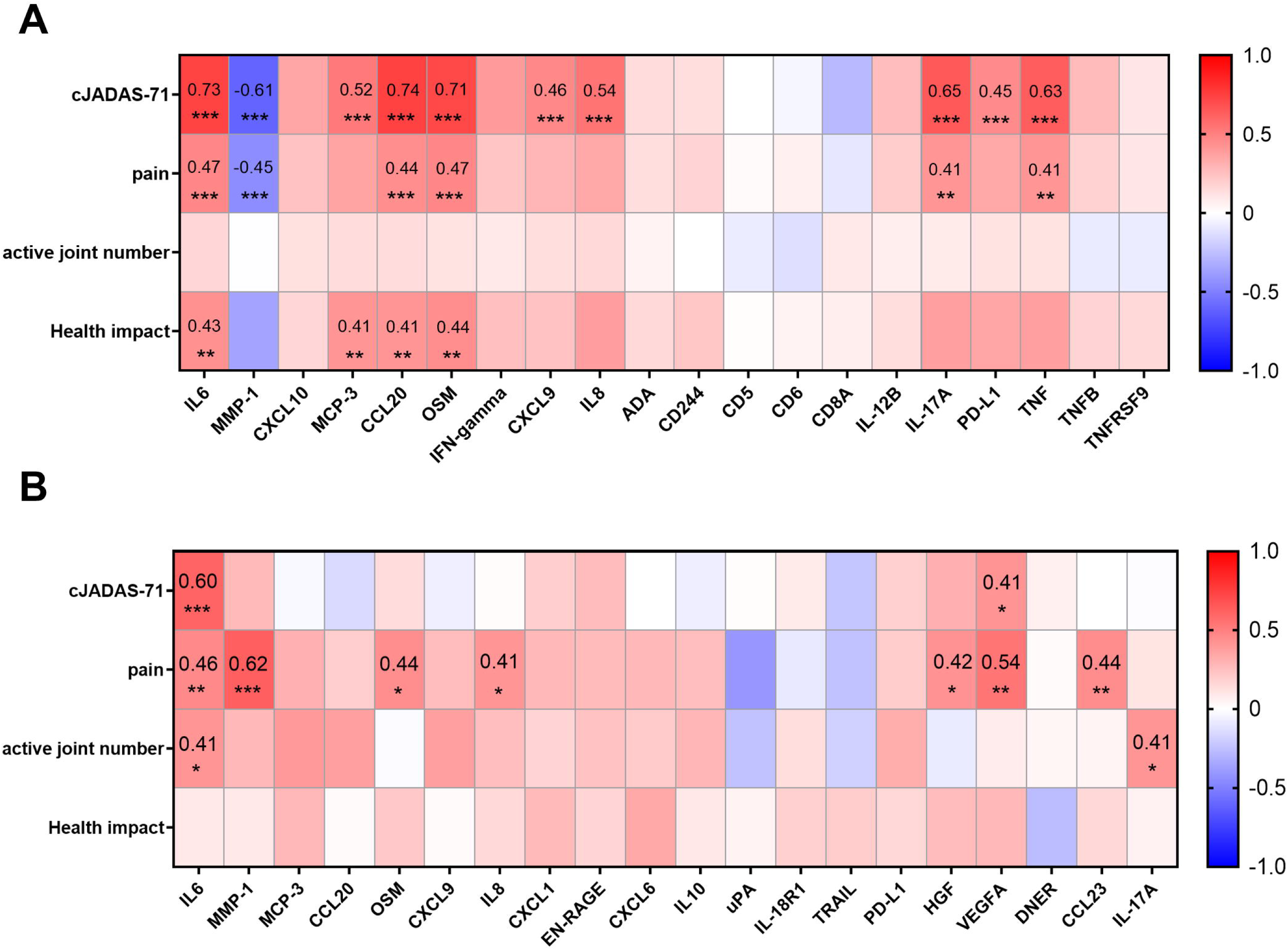
Correlations between DEPs and selected clinical parameters. A) Heatmap visualization of the correlations between DEPs in SF and selected clinical parameters (cJADAS-71, pain, active joint number, and health impact) reveal the proteins related to disease activity, pain, or the score of disease impact on patients’ life. B) Heatmap visualization of the correlations between proteins in plasma and selected clinical parameters. Statistics: Pearson correlation analysis was applied to the correlations between DEPs and cJADAS-71/pain, and Spearman correlation analysis was applied to the correlations between DEPs and active joint number/health impact. * P < 0.05, ** P < 0.01, *** P < 0.001

In plasma, only IL6, and VEGFA correlated with cJADAS-71(Figure 4B, Figure S3A). 8 proteins including MMP-1, VEGFA, and IL6 correlated with pain (Figure 4B, Figure S3A). Notably, plasmatic MMP-1 level showed the highest correlation with pain in oJIA patients (r=0.62, *P*<0.001), suggesting a great potential to serve a biomarker indicating patients’ pain. In addition, IL6, IL-17A and 4E-BP1 showed correlation with active joint number. No protein in plasma was found to correlate with health impact (Figure 4B, Figure S3A).

### 3.6 Inflammation profiles of oJIA patients in the early stage may predict progression of disease

The clinical course and therapeutic outcomes in oJIA patients exhibit substantial variability, encompassing scenarios of complete remission without medication to more severe and prolonged arthritis[18]. Currently, it is difficult to predict progression and therapeutic outcomes in clinical practice. It would be of great value to identify biomarkers that could aid in the prediction of disease progression or treatment outcomes using the inflammation profiles in treatment naïve and early stage oJIA patients.

To investigate prognostic value of the DEPs identified in this study, a 5-year follow-up was conducted for 54 out of the 62 oJIA patients who had SF samples collected at baseline. The disease status was defined from the registry and medical record data, and “Remission” was defined as clinical remission without DMARDs or biological DMARDs and no flare for the past 2 years. “Non-remission” included the patients that needed DMARDs or biological DMARDs or flared in the past 2 years (Figure 5A). We divided the cohort into “remission group” (RG, n=15; 28%) and “non-remission group” (NRG, n=39; 72%) (Figure 5B).

**Figure 5.**
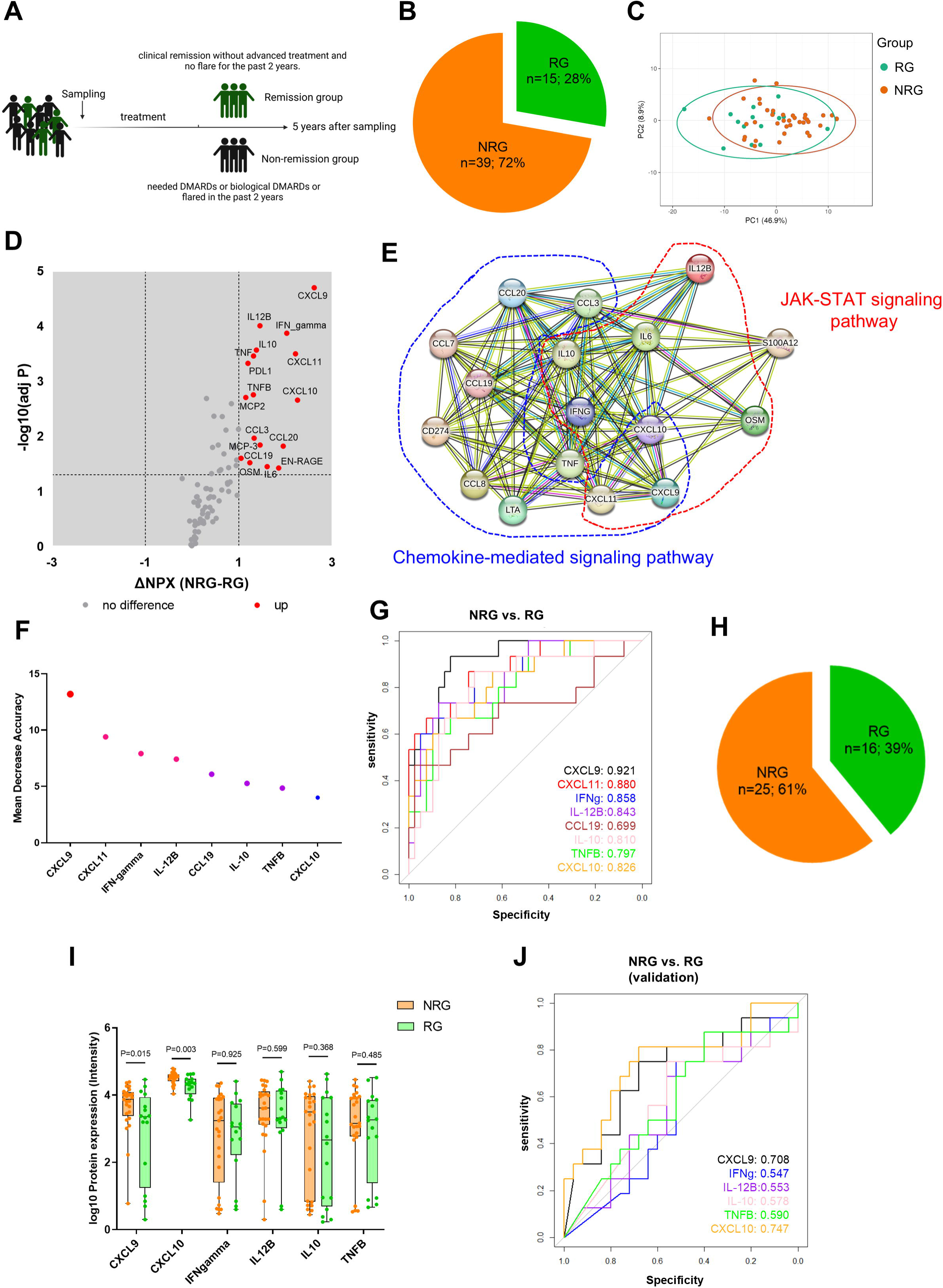
Early inflammation profiles in oJIA predict disease progression. A) Schematic diagram illustrates grouping of oJIA patients based on 5 years follow-up with sampling of synovial fluid at baseline. B) Pie chart presents the percentage of remission group (RG) and non-remission group (NRG) in oJIA patients (n=54). C) PCA results of RG and NRG based on 76 included proteins in SF present an overlap between two groups. Confidence level of the ellipses is 0.95. D-E) Volcano plot show 17 proteins increased in NRG and further STRING analysis highlights the roles of JAK-STAT pathway in NRG. F) Random forest analysis reveals the proteins contributing the separation between NRG and RG. G) ROC analysis was applied to test the ability of each individual protein to separate NRG and RG. H) Pie chart presents the percentage of RG and NRG in validation cohort (n=41). I) Boxplot presents the expression of selected proteins between NRG and RG in validation cohort. J) ROC analysis results in validation cohort reveals CXCL9 and CXCL10 have a good predictive ability to distinguish non-remission individuals.

The baseline concentration in SF showed that 17 proteins including CXCL9, IFN-gamma, CXCL11, IL-12B, and CXCL10 were increased in the NRG, despite a large overlap in profiles between RG and NRG (Figure 5C-5D). STRING analysis showed these increased proteins were related to chemokines and chemokine mediated pathway. Importantly, increased proteins also related with JAK-STAT pathway (Figure 5E). We performed RF analysis and found CXCL9, CXCL11, INF-gamma, and IL-12B were most important in distinguishing NRG in oJIA patients (Figure 5F). Furthermore, ROC analysis results revealed that CXCL9, CXCL11, IFN-gamma, IL-12B, CXCL10, and IL10 had a great capacity to distinguish between NRG and RG, especially CXCL9, with AUC 0.921(95%CI: 0.849-0.993, P<0.001) (Figure 5G, Table S8).

To validate our findings, we used a publicly available independent cohort that encompassed 41 patients with 16 patients in remission group (patients who received no medication or received NSAIDs only during the disease course) and 25 non-remission (patients who need advanced treatment) from Wilmington, USA[16] (Figure 6H). CXCL9, IFN-gamma, IL-12B, CXCL10, IL10, and TNFB were included for analysis, while CXCL11, and CCL19 had to be excluded since they were not part of the Luminex multiplex panel used in the validation cohort. Similar to our data set, CXCL9 and CXCL10 were significantly increased in NRG. No other proteins displayed differences between NRG and RG in the validation cohort (Figure 5I). ROC analysis revealed that CXCL9, and CXCL10 had a good prediction power to distinguish non-remission individual in the oJIA group (Figure 5J, Table S8).

## 4. Discussion

The immunopathogenesis of oJIA is highly complex and not well studied[18–20]. Comprehensive inflammation profiling of early-stage, treatment naive oJIA patients is much needed to gain profound insights into the molecular basis of oJIA pathophysiology and to discover biomarkers for early diagnosis and prognosi.

In this study, we investigated a panel of 92 inflammatory proteins in SF and plasma from oJIA patients and compared them with three different control groups: SF from knee-healthy individuals and from individuals with acute knee injury. oJIA plasma inflammation profiles were compared with plasma from age and sex-matched healthy individuals. When comparing the oJIA cohort with the knee-healthy individuals we found increased levels of 47 proteins in SF, while only two proteins (IL6 and MMP-1) were elevated in plasma when comparing oJIA to healthy controls. These findings suggest that inflammation in oJIA is predominantly confined to the affected joints rather than systemic. Thus, studies using SF rather than plasma samples can provide a more in-depth understanding of oJIA pathogenesis.

Further analysis based on the DEPs in SF, cytokines and chemokines (e.g., IL6, TNF, IFN-gamma, MCP-1, MCP-2, CCL3, and CCL4) suggested the recruitment of immune cells plays an important role in oJIA development and progression[21, 22]. Additionally, many proteins (e.g., ADA, CD5, CD6, and CD8A) related to T cell function indicated activated T cells as being part of the pathogenesis. STRING analysis of SF DEPs identified IL6, TNF, IL10, and IFN-gamma as hub proteins, highlighting their key roles in oJIA pathogenesis. Consistently, increased IL6, TNF, and IFN-gamma have been observed in JIA in many previous studies, and corresponding inhibitors have been applied in clinical practice[23–25]. Interestingly, we found that TNF and IFN-gamma specifically increase in oJIA patients, while IL-6 increases in both oJIA and knee injury patients, suggesting that anti-TNF treatment might more precisely target oJIA compared to IL-6 blockade. IL10, a protein functioning as an anti-inflammatory agent, was increased in SF from oJIA patients. Previous studies also reported higher levels of IL10 in plasma of active systemic JIA patients[26]. The increase in IL-10 is likely a compensatory down-regulation mechanism within the inflammatory response. However, despite this increase, an imbalance persists, reflected by the predominance of elevated pro-inflammatory cytokines and chemokines among the identified DEPs. In addition to these findings, decreased levels of OPG and increased levels of MMP-1, MMP-10, TRANCE and VEGFA in SF reflect the pathological changes associated with bone and cartilage destruction and with pathological angiogenesis in oJIA[27, 28].

Comparison of DEPs between oJIA and knee injury versus controls revealed that oJIA exhibits a distinct inflammation profile from acute inflammation following knee injury. Thirteen proteins (e.g., ADA, CD5, CD6, and CD8A) associated with lymphocytes that were specifically increased in oJIA as compared to controls, were also specifically increased when compared to knee injury patients. Further RF and ROC analysis of oJIA, knee injury, and control groups revealed that ADA, CD8A, and IFN-gamma in SF can serve as diagnostic biomarkers for oJIA. ADA is an important regulatory molecule in the maturation and maintenance of the immune system. It functions by catalyzing the deamination of adenosine, which in turn acts as an immunosuppressive signal, preventing excessive inflammatory responses[29]. Increased ADA has been observed in many autoimmune diseases, such as SLE and RA[29, 30]. However, there are few studies associating ADA with JIA. Lee et al. reported that ADA2 can serve as a biomarker of macrophage activation syndrome (MAS) in sJIA patients[31]. In this study, we observed increased ADA in SF from oJIA patients and further identified it as a new diagnostic biomarker for oJIA. Symons et al. reported higher CD8 levels in serum and SF in RA patients[32]. However, no previous studies have investigated soluble CD8 in oJIA. Increased CD8A in SF from oJIA patients in our study indicated CD8+ T cell activation in oJIA, and we also identified the diagnostic value of CD8A in SF. IFN-gamma has been extensively studied in JIA and identified to play an important role in the pathogenesis of JIA[25, 33]. Consistently, we observed increased levels of IFN-gamma in SF from oJIA patients and, like ADA and CD8A, identified it as a potential diagnostic biomarker for oJIA. However, the lack of correlation between plasma and SF expression of these proteins means they can only serve as diagnostic biomarkers using SF rather than plasma samples. Interestingly, the proteins only increased in knee injury indicated activation of the phosphoinositide 3-kinases (PI3K) pathway. The PI3K pathway has been identified as a crucial pathway that regulates cell survival, differentiation, and growth[34, 35]. Our results thus point to a stronger regenerative activity going on in knee injury patients than in oJIA patients.

In line with the higher number of DEPs in oJIA SF, a greater number of correlations between protein levels and available clinical parameters compared to those in plasma could be revealed. In SF, 10 proteins were found to have moderate or strong and highly significant correlations with cJADAS-71. Notably, CCL20 exhibited the strongest correlation with cJADAS-71, suggesting that it can serve as a highly effective biomarker for disease activity in clinical assessments. Higher levels of CCL20 have been observed in RA and JIA patients [36, 37]. CCL20/CCR6 signaling is known to inhibit Treg cell differentiation while promoting Th17 cell differentiation[38]. The high correlation between CCL20 and disease activity suggests a dysregulated Th17/Treg balance in the immunopathogenesis of oJIA, and that CCL20 would be a therapeutic target. Unexpectedly, MMP-1 negatively correlated with both cJADAS-71 and pain, and a negative correlation was identified between its expression in plasma and SF. Further investigation is needed to better understand the cause of this relationship. In addition, 6 proteins showed significant correlation with pain, and 4 proteins significantly correlated with health impact, although with weak to moderate strength. The defined proteins could be of potential value for assessing patient quality of life in clinical settings and deserves further studies on larger number of samples.

Although a higher number of significant correlations could be defined for SF, some plasma proteins also showed significant correlations with these clinical parameters. Similar to SF IL6, plasma IL6 showed a good correlation with cJADAS-71. Thus, plasma IL6 would be a viable biomarker option in clinical practice for disease activity when SF samples are not available. Plasma MMP-1 showed the highest correlation with pain. Quantitative biomarkers for pain are lacking in clinical practice today. Our data on MMP-1 implies that MMP-1 might fulfill this clinical need, and further studies are warranted. VEGFA, a cytokine associated with angiogenesis, demonstrated moderate significant correlations with both cJADAS-71 and pain when measured in plasma and should be explored further.

For 54 of the included oJIA patients, clinical outcome data were recorded at a time point 5 years after SF sampling. This provided the possibility to analyze whether inflammation profiles in the early, treatment naïve, stage of disease could predict disease remission after 5 years. We identified 17 DEPs in SF of non-remission patients and ensuing STRING analysis showed enrichment in the JAK-STAT pathway, besides the chemokine-mediated pathways previously indicated in the whole cohort. The defined DEPs CXCL9, CXCL10, and CXCL11 are induced by the DEP IFN-gamma via the JAK-STAT pathway. By performing ROC-analyses we revealed high predictive biomarker features of eight DEPs, with CXCL9 and CXCL11 being the two top biomarkers. IFN-gamma along with its induced cytokines have been reported as biomarkers of macrophage activation syndrome in systemic JIA patients [31, 39]. However, few studies have investigated their potential as biomarkers in oJIA. Al-Jaberi et al reported in a longitudinal study, that early increased levels of CXCL9 and CXCL10 were associated with later need of advanced therapy[16]. In our study, we further demonstrated the predictive potential of CXCL9 and CXCL10 for disease progression in oJIA. Our data suggests that oJIA patients in the early stage with high levels of CXCL9 and CXCL10 have a higher risk of not achieving remission and could benefit from an early application of JAKi, though JAK inhibitors (JAKi) are not first-line treatment according to current clinical practice[1]. Further research is needed to fully establish the role of CXCL11 in oligoarticular arthritis and its utility as a biomarker in this JIA subtype.

Strengths of our study are the parallel analysis of SF and plasma oJIA samples and the selection of treatment naïve patients with short disease duration and well-defined clinical characteristics for the cross-sectional analysis. The longitudinal data on disease outcomes allowing the study of prognostic biomarkers adds to its strengths. Comparative analyses of SF samples from non-arthritic controls are rare and adds to this study’s novelty. There are also limitations in this study. The limited sample size of our cohort constrained the validation of our findings within this study population. The SF control and knee-injury cohorts are not sex- and age-matched with the oJIA group. Additionally, the reference plasma samples and reference SF samples are not from the same individuals, which may lead to differences in the standard for comparison.

In conclusion, by inflammation profiling SF samples from oJIA and control cohorts, we have identified activated molecular pathways and biomarkers specific for oJIA. In contrast, plasma inflammation profiles were similar between oJIA and controls. Our data implicate that biomarkers associated with T cell activation show promise for diagnostic purposes while proteins connected to an activation of the JAK/STAT signalling pathway show promise as predictive biomarkers for a chronic disease progression and could aid in patient stratification and therapy decisions. While SF analysis revealed more biomarkers correlating with clinical parameters than plasma analysis, it is noteworthy that increased plasma levels of IL6 and MMP-1 correlated with disease activity and pain. However, to further deepen the knowledge of oJIA immunopathogenesis and to establish implementation of new diagnostic and prognostic biomarkers in the clinics, validation of our results in independent cohorts is warranted.

