## Supplementary tables and figures for "Inflammation-profiling reveals activated pathways and biomarkers with predictive potential in oligoarticular JIA"

**Supplementary table 1. Demographics and disease characteristics of the population in the validation cohort**

|  | oJIA patients for validation  (SF) |
| --- | --- |
| Sample size(n) | 41 |
| Sex(F) | 34  *P*=0.979 (vs oJIA SF in table 1) |
| Age at sampling (median/range) | NA* |
| **RF** Measurement(n) | 28 |
| positive | 1 (3.57%) |
| **ANA** Measurement(n) | 35 |
| positive | 25 (71.43%) |
| **HLA-B27** Measurement(n) | 22 |
| positive | 4 (18.18%) |

*: The information is not clearly stated in the study involving the validation cohort. Details can be referenced in PMID: 36342195.

**Supplementary table 2. The 92 proteins included in Olink’s Target Inflammation panel**

| UniProt ID | Proteins (Olink ID) | Full name | Call rate* in plasma | Call rate in SF |
| --- | --- | --- | --- | --- |
| P00813 | ADA | adenosine deaminase | >80% | >80% |
| Q5T4W7 | ARTN | artemin | 20-80% | <20% |
| O15169 | AXIN1 | axin 1 | >80% | 20-80% |
| Q14790 | CASP8 | caspase 8 | 20-80% | >80% |
| P51671 | CCL11 | C-C motif chemokine ligand 11 | >80% | >80% |
| Q99616 | CCL13 (MCP-4) | C-C motif chemokine ligand 13 | >80% | >80% |
| Q99731 | CCL19 | C-C motif chemokine ligand 19 | >80% | >80% |
| P13500 | CCL2 (MCP-1) | C-C motif chemokine ligand 2 | >80% | >80% |
| P78556 | CCL20 | C-C motif chemokine ligand 20 | >80% | >80% |
| P55773 | CCL23 | C-C motif chemokine ligand 23 | >80% | >80% |
| O15444 | CCL25 | C-C motif chemokine ligand 25 | >80% | 20-80% |
| Q9NRJ3 | CCL28 | C-C motif chemokine ligand 28 | >80% | >80% |
| P10147 | CCL3 | C-C motif chemokine ligand 3 | >80% | >80% |
| P13236 | CCL4 | C-C motif chemokine ligand 4 | >80% | >80% |
| P80098 | CCL7 (MCP-3) | C-C motif chemokine ligand 7 | 20-80% | 20-80% |
| P80075 | CCL8 (MCP-2) | C-C motif chemokine ligand 8 | >80% | >80% |
| Q9BZW8 | CD244 | CD244 molecule | >80% | >80% |
| Q9NZQ7 | CD274 (PD-L1) | CD274 molecule | >80% | >80% |
| P25942 | CD40 | CD40 molecule | >80% | >80% |
| P06127 | CD5 | CD5 molecule | >80% | >80% |
| P30203 | CD6 | CD6 molecule | >80% | >80% |
| P01732 | CD8A | CD8a molecule | >80% | >80% |
| Q9H5V8 | CDCP1 | CUB domain containing protein 1 | >80% | >80% |
| P21583 | KITLG (SCF) | c-Kit ligand | >80% | >80% |
| P09603 | CSF1 | colony stimulating factor 1 | >80% | >80% |
| P28325 | CST5 | cystatin D | >80% | 20-80% |
| P78423 | CX3CL1 | C-X3-C motif chemokine ligand 1 | >80% | 20-80% |
| P09341 | CXCL1 | C-X-C motif chemokine ligand 1 | >80% | >80% |
| P02778 | CXCL10 | C-X-C motif chemokine ligand 10 | >80% | >80% |
| O14625 | CXCL11 | C-X-C motif chemokine ligand 11 | >80% | >80% |
| P42830 | CXCL5 | C-X-C motif chemokine ligand 5 | >80% | 20-80% |
| P80162 | CXCL6 | C-X-C motif chemokine ligand 6 | >80% | >80% |
| Q07325 | CXCL9 | C-X-C motif chemokine ligand 9 | >80% | >80% |
| Q8NFT8 | DNER | delta/notch like EGF repeat containing | >80% | >80% |
| Q13541 | EIF4EBP1 (4E-BP1) | eukaryotic translation initiation factor 4E binding protein 1 | >80% | >80% |
| O95750 | FGF19 | fibroblast growth factor 19 | >80% | >80% |
| Q9NSA1 | FGF21 | fibroblast growth factor 21 | 20-80% | 20-80% |
| Q9GZV9 | FGF23 | fibroblast growth factor 23 | 20-80% | 20-80% |
| P12034 | FGF5 | fibroblast growth factor 5 | 20-80% | <20% |
| P49771 | FLT3LG (Flt3L) | fms related tyrosine kinase 3 ligand | >80% | >80% |
| P39905 | GDNF | glial cell derived neurotrophic factor | 20-80% | <20% |
| P14210 | HGF | hepatocyte growth factor | >80% | >80% |
| P22301 | IL10 | interleukin 10 | >80% | 20-80% |
| Q13651 | IL10RA | interleukin 10 receptor subunit alpha | 20-80% | 20-80% |
| Q08334 | IL10RB | interleukin 10 receptor subunit beta | >80% | >80% |
| P29460 | IL12B | interleukin 12B | >80% | >80% |
| P35225 | IL13 | interleukin 13 | <20% | <20% |
| Q13261 | IL15RA | interleukin 15 receptor subunit alpha | 20-80% | 20-80% |
| Q16552 | IL17A | interleukin 17A | 20-80% | 20-80% |
| Q9P0M4 | IL17C | interleukin 17C | 20-80% | 20-80% |
| Q14116 | IL18 | interleukin 18 | >80% | >80% |
| Q13478 | IL18R1 | interleukin 18 receptor 1 | >80% | >80% |
| P01583 | IL1A | interleukin 1 alpha | <20% | <20% |
| P60568 | IL2 | interleukin 2 | <20% | <20% |
| Q9NYY1 | IL20 | interleukin 20 | <20% | <20% |
| Q9UHF4 | IL20RA | interleukin 20 receptor subunit alpha | 20-80% | <20% |
| Q8N6P7 | IL22RA1 | interleukin 22 receptor subunit alpha 1 | 20-80% | <20% |
| Q13007 | IL24 | interleukin 24 | 20-80% | 20-80% |
| P14784 | IL2RB | interleukin 2 receptor subunit beta | 20-80% | <20% |
| O95760 | IL33 | interleukin 33 | <20% | <20% |
| P05112 | IL4 | interleukin 4 | 20-80% | <20% |
| P05113 | IL5 | interleukin 5 | 20-80% | <20% |
| P05231 | IL6 | interleukin 6 | >80% | >80% |
| P13232 | IL7 | interleukin 7 | >80% | 20-80% |
| P10145 | IL8 | C-X-C motif chemokine ligand 8 | >80% | >80% |
| P01579 | IFN-gamma | Interferon gamma | **#** | 20-80% |
| P15018 | LIF | LIF interleukin 6 family cytokine | <20% | 20-80% |
| P42702 | LIFR | LIF receptor subunit alpha | >80% | 20-80% |
| P03956 | MMP-1 | matrix metallopeptidase 1 | >80% | >80% |
| P09238 | MMP-10 | matrix metallopeptidase 10 | >80% | >80% |
| P01138 | NGF (Beta-NGF) | nerve growth factor | 20-80% | 20-80% |
| Q99748 | NRTN | neurturin | <20% | <20% |
| O00300 | OPG | TNF receptor superfamily member 11b | >80% | >80% |
| P13725 | OSM | oncostatin M | >80% | >80% |
| P00749 | PLAU (uPA) | Urolinase-type plasminoen activator | >80% | >80% |
| P80511 | S100A12 (EN-RAGE) | S100 calcium binding protein A12 | >80% | 20-80% |
| Q8IXJ6 | SIRT2 | sirtuin 2 | 20-80% | 20-80% |
| Q13291 | SLAMF1 | signaling lymphocytic activation molecule family member 1 | 20-80% | 20-80% |
| P20783 | NTF3 (NT-3) | Neurotrophin 3 | >80% | <20% |
| O95630 | STAMBP | STAM binding protein | >80% | >80% |
| P50225 | SULT1A1 (ST1A1) | sulfotransferase family 1A member 1 | 20-80% | >80% |
| P01135 | TGFA | transforming growth factor alpha | >80% | >80% |
| P01137 | TGFB1 (LAP TGF-beta-1) | transforming growth factor beta 1 | >80% | >80% |
| P01375 | TNF | Tumor necrosis factor | **#** | >80% |
| P01374 | TNFB | lymphotoxin alpha | >80% | >80% |
| Q07011 | TNFRSF9 | TNF receptor superfamily member 9 | >80% | >80% |
| P50591 | TNFSF10 (TRAIL) | TNF superfamily member 10 | >80% | >80% |
| O14788 | TNFSF11 (TRANCE) | TNF superfamily member 11 | >80% | 20-80% |
| O43508 | TNFSF12 (TEWAK) | TNF superfamily member 12 | >80% | >80% |
| O43557 | TNFSF14 | TNF superfamily member 14 | >80% | >80% |
| Q969D9 | TSLP | thymic stromal lymphopoietin | <20% | <20% |
| P15692 | VEGFA | vascular endothelial growth factor A | >80% | >80% |

* Call rate: The percentage of protein expression above the limit of detection (LOD).

\# TNF and IFN-gamma in plasma had to be excluded from the analysis due to a change in antibodies between the two runs.

**Supplementary table 3. Comparison of protein expression between oJIA and HC in plasma.**

| Proteins | ΔNPX(oJIA-HC)/Fold Change | Padj |
| --- | --- | --- |
| IL8 | -0.45495 | 0.00731 |
| VEGFA | -0.31008 | 0.010065 |
| MCP3 | 0.412288 | 0.007017 |
| GDNF | 0.158603 | 0.096658 |
| CDCP1 | 0.082333 | 0.292955 |
| CD244 | -0.4571 | 1.53E-05 |
| IL7 | -1.23142 | 1.24E-08 |
| OPG | -0.18866 | 0.005863 |
| LAP TGF beta1 | -0.46318 | 0.00536 |
| uPA | -0.3359 | 1.48E-06 |
| IL6 | 1.121835 | 1.46E-06 |
| IL-17C | -0.28777 | 0.020603 |
| MCP1 | -0.09696 | 0.262453 |
| IL-17A | -0.14829 | 0.231815 |
| CXCL11 | -0.85893 | 0.000565 |
| AXIN1 | -1.83199 | 8.33E-07 |
| TRAIL | 0.148263 | 0.012288 |
| IL20RA | -0.13835 | 0.220061 |
| CXCL9 | 0.715639 | 0.000282 |
| CST5 | -0.05528 | 0.533049 |
| IL2RB | -0.13226 | 0.507498 |
| OSM | 0.391239 | 0.098584 |
| CXCL1 | -1.51166 | 2.56E-09 |
| CCL4 | -0.60085 | 0.000132 |
| CD6 | -0.57416 | 0.000534 |
| SCF | -0.50277 | 7.17E-06 |
| IL18 | -0.47823 | 0.003541 |
| SLAMF1 | -0.11975 | 0.10288 |
| TGF alpha | 0.088854 | 0.377377 |
| MCP4 | -0.95978 | 0.000103 |
| CCL11 | -0.22883 | 0.028985 |
| TNFSF14 | -0.51477 | 0.004187 |
| FGF23 | -0.04415 | 0.506034 |
| IL10RA | 0.124838 | 0.445642 |
| FGF5 | 0.035839 | 0.402355 |
| MMP-1 | 1.616392 | 4.81E-08 |
| LIF-R | -0.11054 | 0.029624 |
| FGF21 | 0.03075 | 0.908151 |
| CCL19 | 0.130362 | 0.369396 |
| IL15RA | -0.15697 | 0.004749 |
| IL10RB | -0.14595 | 0.014073 |
| IL22 RA1 | -0.1431 | 0.148369 |
| IL18R1 | 0.042061 | 0.567235 |
| PD-L1 | -0.10389 | 0.169885 |
| beta NGF | 0.104115 | 0.000147 |
| CXCL5 | -2.18439 | 1.81E-09 |
| TRANCE | -0.43413 | 0.005202 |
| HGF | -0.09654 | 0.286226 |
| IL-12B | 0.132002 | 0.271254 |
| IL24 | 0.106845 | 0.431216 |
| ARTN | -0.01998 | 0.874857 |
| MMP10 | -0.16374 | 0.320137 |
| IL10 | 0.193946 | 0.156282 |
| CCL23 | -0.039 | 0.736538 |
| CD5 | -0.45901 | 1.39E-05 |
| CCL3 | -0.37022 | 0.005371 |
| FIt3L | -0.2028 | 0.01619 |
| CXCL6 | -2.02733 | 1.01E-12 |
| CXCL10 | 0.199601 | 0.218196 |
| 4E-BP1 | -1.27332 | 0.000672 |
| SIRT2 | -2.4673 | 3.41E-09 |
| CCL28 | -0.08982 | 0.492182 |
| EN-RAGE | 0.4381 | 0.087958 |
| CD40 | -0.42336 | 0.000334 |
| FGF19 | -0.33478 | 0.123523 |
| IL4 | -0.38773 | 0.014238 |
| MCP-2 | -0.79872 | 5.9E-05 |
| CASP-8 | -1.72469 | 6.56E-09 |
| CCL25 | 0.065966 | 0.595991 |
| CX3CL1 | -0.17421 | 0.057037 |
| TNFRSF9 | -0.0924 | 0.284155 |
| NT3 | -0.29478 | 0.004337 |
| TWEAK | -0.37944 | 5.58E-06 |
| CCL20 | -0.12224 | 0.440346 |
| ST1A1 | -2.21836 | 9.76E-12 |
| STAMBP | -1.5353 | 1.71E-06 |
| IL5 | 0.130919 | 0.620563 |
| ADA | -0.58323 | 5.49E-06 |
| TNFB | -0.18352 | 0.069674 |
| CSF-1 | 0.174194 | 7.08E-05 |
| DNER | -0.10847 | 0.038765 |
| CD8A | 0.05124 | 0.735633 |

**Supplementary table 4. Comparison of protein expression between oJIA and controls in SF.**

| Proteins | ΔNPX(oJIA-Ctrl)/Fold Change | P_adj_ |
| --- | --- | --- |
| IL8 | 4.698336 | 9.13026E-08 |
| VEGFA | 1.735228 | 2.24533E-05 |
| CD8A | 2.756263 | 0.007427318 |
| MCP-3 | 5.568467 | 9.14715E-09 |
| CDCP1 | 2.966906 | 3.32618E-06 |
| CD244 | 1.021684 | 0.008381833 |
| IL7 | -0.16748 | 0.194980419 |
| OPG | -1.05865 | 0.001271686 |
| LAP TGF-beta-1 | 0.669239 | 0.084966749 |
| uPA | 2.73788 | 3.53478E-08 |
| IL6 | 9.936788 | 2.45878E-13 |
| IL17-C | -0.2565 | 0.00251788 |
| MCP-1 | 1.805311 | 5.35244E-06 |
| IL-17A | 1.074858 | 2.23812E-08 |
| CXCL11 | 3.792661 | 0.000104435 |
| AXIN1 | 0.722744 | 0.036454152 |
| TRAIL | 2.613372 | 4.77689E-06 |
| CXCL9 | 4.898232 | 1.86433E-06 |
| CST5 | -0.48644 | 0.075379283 |
| OSM | 5.274957 | 4.49449E-10 |
| CXCL1 | 3.80883 | 1.08274E-06 |
| CCL4 | 2.152332 | 0.000122108 |
| CD6 | 1.937621 | 0.001919013 |
| SCF | 0.446005 | 0.216543424 |
| IL18 | 1.485932 | 0.000347004 |
| SLAMF1 | 0.944745 | 0.000385535 |
| TGF-alpha | 0.122191 | 0.506557792 |
| MCP-4 | 2.569724 | 0.000145187 |
| CCL11 | 0.860265 | 0.00764207 |
| TNFSF14 | 2.426626 | 0.000369709 |
| FGF-23 | -0.42007 | 0.003854863 |
| IL-10RA | 0.273019 | 0.002181263 |
| MMP-1 | 7.253249 | 1.02633E-05 |
| LIF-R | 0.610182 | 6.75136E-05 |
| FGF-21 | -0.09121 | 0.68169169 |
| CCL19 | 3.35161 | 2.72672E-05 |
| IL-15RA | 0.491745 | 0.001194676 |
| IL-10RB | 0.544405 | 0.041227975 |
| IL-18R1 | 2.668828 | 1.82396E-05 |
| PD-L1 | 2.317713 | 6.30662E-06 |
| Beta-NGF | 0.329453 | 0.000175599 |
| CXCL5 | 0.344977 | 0.651085465 |
| TRANCE | 3.457566 | 1.21399E-08 |
| HGF | 1.988508 | 1.17709E-05 |
| IL-12B | 3.225927 | 1.30639E-05 |
| IL-24 | 0.626764 | 0.000269149 |
| MMP-10 | 1.965739 | 1.94527E-05 |
| IL10 | 2.840577 | 3.64227E-10 |
| TNF | 1.932992 | 1.14209E-12 |
| CCL23 | 1.337726 | 0.000325912 |
| CD5 | 2.827946 | 0.001442773 |
| CCL3 | 3.347924 | 6.84247E-07 |
| Flt3L | -0.81144 | 0.011070519 |
| CXCL6 | 2.91996 | 6.67037E-05 |
| CXCL10 | 5.938753 | 2.53035E-06 |
| 4E-BP1 | 1.504306 | 0.043729046 |
| SIRT2 | 1.2705 | 0.027586716 |
| CCL28 | -0.11708 | 0.227874118 |
| DNER | 1.642512 | 0.000171696 |
| EN-RAGE | 3.706395 | 6.71837E-08 |
| CD40 | 1.860006 | 0.000136123 |
| IFN-gamma | 5.036109 | 1.32863E-06 |
| FGF-19 | 0.954897 | 0.002637795 |
| LIF | 3.368759 | 6.13635E-15 |
| MCP-2 | 2.532883 | 8.35624E-06 |
| CASP-8 | -0.00403 | 0.993564349 |
| CCL25 | 0.576033 | 0.047117474 |
| CX3CL1 | 0.539391 | 0.072609258 |
| TNFRSF9 | 1.77899 | 0.000682607 |
| TWEAK | 0.332614 | 0.141401169 |
| CCL20 | 5.448214 | 1.75979E-15 |
| ST1A1 | 0.377272 | 0.39632731 |
| STAMBP | 0.619321 | 0.187272928 |
| ADA | 4.05637 | 2.11534E-07 |
| TNFB | 2.581661 | 2.80474E-05 |
| CSF-1 | 0.339867 | 0.259869912 |

**Supplementary table 5. ROC analysis results of selected markers distinguish oJIA from juvenile knee injury and controls.**

| Proteins | AUC | 95% CI | P |
| --- | --- | --- | --- |
| ADA | 0.996 | 0.990-1 | 1.03E-16 |
| CD8A | 0.947 | 0.890-1 | 7.16E-14 |
| IFN-gamma | 0.960 | 0.917-1 | 1.46E-14 |
| IL6 | 0.565 | 0.434-0.697 | 0.274126 |
| LIF | 0.571 | 0.430-0.712 | 0.232857 |
| CXCL10 | 0.863 | 0.787-0.939 | 1.27E-09 |
| CASP-8 | 0.807 | 0.706-0.908 | 2.83E-07 |
| CXCL9 | 0.868 | 0.787-0.949 | 7.42E-10 |
| IL-12B | 0.855 | 0.754-0.955 | 2.91E-09 |
| MCP-3 | 0.686 | 0.570-0.801 | 0.001868 |
| OPG | 0.790 | 0.679-0.901 | 1.23E-06 |
| MMP-1 | 0.755 | 0.629-0.880 | 2.04E-05 |

**Supplementary table 6. Correlations of DEPs between plasma and SF (ranked by r)**

| DEPs | r | P value |
| --- | --- | --- |
| IL6 | 0.621453 | 4.04E-05 |
| MMP-10 | 0.541508 | 0.000538 |
| EN-RAGE | 0.53321 | 0.000679 |
| MMP-1 | -0.48237 | 0.002500 |
| VEGFA | 0.462107 | 0.003985 |
| CCL4 | 0.460585 | 0.004122 |
| 4E-BP1 | 0.436924 | 0.006853 |
| TNFSF14 | 0.37548 | 0.022017 |
| DNER | 0.372978 | 0.02299 |
| CD8A | 0.366027 | 0.025881 |
| CCL19 | 0.357778 | 0.029696 |
| IL-12B | 0.344732 | 0.036665 |
| CXCL1 | 0.318891 | 0.054389 |
| MCP-2 | 0.318528 | 0.05468 |
| CD6 | 0.315602 | 0.057068 |
| uPA | 0.309414 | 0.06239 |
| CD40 | 0.298901 | 0.072324 |
| AXIN1 | 0.295049 | 0.076259 |
| HGF | 0.276811 | 0.097202 |
| CCL3 | 0.266783 | 0.110453 |
| CXCL10 | 0.253208 | 0.130508 |
| MCP-3 | 0.246228 | 0.141815 |
| CXCL11 | 0.23546 | 0.160646 |
| IL-18R1 | 0.233638 | 0.164002 |
| IL8 | 0.23025 | 0.170379 |
| CXCL9 | 0.215883 | 0.199391 |
| CD244 | 0.214322 | 0.202739 |
| TRANCE | -0.17521 | 0.299627 |
| CD5 | 0.152091 | 0.368853 |
| SIRT2 | 0.148249 | 0.381211 |
| IL18 | 0.139115 | 0.411553 |
| OSM | 0.135222 | 0.424888 |
| CXCL6 | 0.10963 | 0.518333 |
| PD-L1 | 0.089432 | 0.598624 |
| OPG | -0.06896 | 0.685085 |
| IL10 | 0.055485 | 0.744294 |
| CDCP1 | 0.052258 | 0.758713 |
| TNFRSF9 | 0.051025 | 0.76424 |
| MCP-4 | 0.048559 | 0.775337 |
| IL-17A | -0.0396 | 0.815998 |
| ADA | 0.039554 | 0.816203 |
| TNFB | 0.033667 | 0.843189 |
| MCP-1 | 0.032466 | 0.848717 |
| CCL20 | -0.02859 | 0.8666 |
| CCL23 | 0.016722 | 0.921749 |

**Supplementary table 7. Correlations between DEPs in SF and selected clinical parameters** (significant correlations are marked in red)

| DEPs | cJADAS-71 | | VAS Pain | | Health impact | | Active joint count | |
| --- | --- | --- | --- | --- | --- | --- | --- | --- |
|  | r | p | r | p | Spearman r | P | Spearman r | P |
| IL6 | 0.7258 | <0.0001 | 0.4729 | 0.0003 | 0.4295 | 0.0012 | 0.1639 | 0.2362 |
| MMP-1 | -0.6074 | <0.0001 | -0.4461 | 0.0007 | -0.3588 | 0.0077 | -0.006949 | 0.9602 |
| CXCL10 | 0.3522 | 0.009 | 0.2374 | 0.0839 | 0.1564 | 0.2586 | 0.1175 | 0.3975 |
| MCP-3 | 0.5195 | <0.0001 | 0.3612 | 0.0073 | 0.4102 | 0.0021 | 0.1397 | 0.3138 |
| CCL20 | 0.7405 | <0.0001 | 0.4363 | 0.001 | 0.4128 | 0.0019 | 0.1441 | 0.2984 |
| OSM | 0.71 | <0.0001 | 0.4702 | 0.0003 | 0.4354 | 0.001 | 0.1065 | 0.4435 |
| IFN-gamma | 0.3853 | 0.004 | 0.2276 | 0.0979 | 0.2498 | 0.0685 | 0.09193 | 0.5085 |
| CXCL9 | 0.4634 | 0.0004 | 0.2921 | 0.0321 | 0.235 | 0.0871 | 0.1307 | 0.346 |
| IL8 | 0.5378 | <0.0001 | 0.3392 | 0.0121 | 0.3783 | 0.0048 | 0.1514 | 0.2743 |
| ADA | 0.1442 | 0.2981 | 0.1326 | 0.3393 | 0.1548 | 0.2638 | 0.05227 | 0.7074 |
| CXCL1 | 0.6132 | <0.0001 | 0.3685 | 0.0061 | 0.3755 | 0.0051 | 0.1841 | 0.1828 |
| CXCL11 | 0.3375 | 0.0126 | 0.2368 | 0.0847 | 0.1786 | 0.1964 | 0.1374 | 0.3216 |
| EN-RAGE | 0.6853 | <0.0001 | 0.3748 | 0.0052 | 0.3973 | 0.0029 | -0.002581 | 0.9852 |
| TRANCE | 0.4445 | 0.0008 | 0.2646 | 0.0531 | 0.355 | 0.0084 | 0.005907 | 0.9662 |
| LIF | 0.6351 | <0.0001 | 0.4782 | 0.0003 | 0.3442 | 0.0108 | 0.1751 | 0.2053 |
| CCL19 | 0.2361 | 0.0857 | 0.1828 | 0.1859 | 0.1284 | 0.3547 | 0.003425 | 0.9804 |
| CCL3 | 0.4795 | 0.0002 | 0.2749 | 0.0442 | 0.359 | 0.0077 | 0.1602 | 0.2471 |
| IL-12B | 0.2564 | 0.0612 | 0.2006 | 0.1459 | 0.1261 | 0.3637 | 0.07366 | 0.5966 |
| CDCP1 | 0.3735 | 0.0054 | 0.381 | 0.0045 | 0.2475 | 0.0711 | 0.1108 | 0.4251 |
| CXCL6 | 0.6283 | <0.0001 | 0.3375 | 0.0126 | 0.3411 | 0.0116 | 0.2553 | 0.0625 |
| IL10 | 0.6151 | <0.0001 | 0.4377 | 0.0009 | 0.3287 | 0.0152 | 0.07445 | 0.5926 |
| CD5 | -0.00463 | 0.9735 | 0.02254 | 0.8715 | 0.008579 | 0.9509 | -0.08111 | 0.5599 |
| CD8A | -0.2815 | 0.0392 | -0.1033 | 0.4573 | 0.06616 | 0.6346 | 0.09242 | 0.5062 |
| uPA | 0.5723 | <0.0001 | 0.3978 | 0.0029 | 0.3566 | 0.0081 | 0.09163 | 0.5099 |
| IL-18R1 | 0.5756 | <0.0001 | 0.3439 | 0.0109 | 0.4285 | 0.0012 | 0.09664 | 0.4869 |
| TRAIL | 0.5909 | <0.0001 | 0.3575 | 0.008 | 0.4555 | 0.0005 | 0.05604 | 0.6873 |
| TNFB | 0.2653 | 0.0525 | 0.1848 | 0.1809 | 0.169 | 0.222 | -0.09103 | 0.5127 |
| MCP-4 | 0.07398 | 0.595 | 0.0983 | 0.4795 | 0.07195 | 0.6051 | -0.1383 | 0.3187 |
| MCP-2 | 0.3513 | 0.0092 | 0.2951 | 0.0303 | 0.2811 | 0.0395 | 0.08349 | 0.5484 |
| TNFSF14 | 0.3322 | 0.0141 | 0.1756 | 0.2041 | 0.3317 | 0.0143 | 0.07535 | 0.5882 |
| PD-L1 | 0.4523 | 0.0006 | 0.3403 | 0.0118 | 0.3457 | 0.0104 | 0.1109 | 0.4247 |
| CCL4 | 0.3482 | 0.0099 | 0.1449 | 0.2958 | 0.3029 | 0.026 | 0.04696 | 0.736 |
| HGF | 0.6453 | <0.0001 | 0.4248 | 0.0014 | 0.3561 | 0.0082 | 0.151 | 0.2756 |
| MMP-10 | 0.3715 | 0.0057 | 0.3386 | 0.0123 | 0.2581 | 0.0595 | 0.2812 | 0.0394 |
| CD6 | -0.03639 | 0.7939 | 0.05907 | 0.6714 | 0.05182 | 0.7098 | -0.1297 | 0.3501 |
| TNF | 0.6251 | <0.0001 | 0.4099 | 0.0021 | 0.365 | 0.0067 | 0.1082 | 0.4363 |
| CD40 | 0.3562 | 0.0082 | 0.2155 | 0.1177 | 0.2834 | 0.0378 | 0.06433 | 0.644 |
| MCP-1 | 0.3126 | 0.0214 | 0.3242 | 0.0168 | 0.3324 | 0.0141 | 0.1314 | 0.3434 |
| TNFRSF9 | 0.1049 | 0.4501 | 0.09647 | 0.4877 | 0.1464 | 0.2909 | -0.0822 | 0.5546 |
| VEGFA | 0.5066 | <0.0001 | 0.4263 | 0.0013 | 0.2249 | 0.102 | 0.1208 | 0.3842 |
| DNER | 0.534 | <0.0001 | 0.4928 | 0.0002 | 0.2173 | 0.1145 | 0.1254 | 0.3663 |
| 4E-BP1 | 0.2269 | 0.0989 | 0.2004 | 0.1463 | 0.1506 | 0.277 | 0.2309 | 0.093 |
| IL18 | 0.2719 | 0.0467 | 0.3719 | 0.0056 | 0.2202 | 0.1097 | 0.144 | 0.2987 |
| CCL23 | 0.5237 | <0.0001 | 0.4358 | 0.001 | 0.2947 | 0.0305 | -0.014 | 0.92 |
| SIRT2 | 0.3798 | 0.0046 | 0.3082 | 0.0234 | 0.2664 | 0.0515 | 0.1713 | 0.2154 |
| IL-17A | 0.6546 | <0.0001 | 0.4074 | 0.0022 | 0.366 | 0.0065 | 0.07599 | 0.585 |
| CD244 | 0.1267 | 0.3613 | 0.1656 | 0.2315 | 0.2186 | 0.1123 | -0.000447 | 0.9974 |
| OPG | 0.3721 | 0.0056 | 0.3003 | 0.0274 | 0.1486 | 0.2834 | 0.2218 | 0.107 |

**Supplementary table 8. ROC analysis results of selected markers in our cohort and validation cohort.**

| Proteins | Discovery cohort | | | Validation cohort | | |
| --- | --- | --- | --- | --- | --- | --- |
|  | AUC | 95% CI | P | AUC | 95% CI | P |
| CXCL9 | 0.921 | 0.849-0.993 | 1.9E-6 | 0.708 | 0.538-0.877 | 0.027 |
| CXCL11 | 0.880 | 0.779-0.982 | 1.7E-5 |  |  |  |
| IFN-gamma | 0.858 | 0.752-0.965 | 5.2E-5 | 0.548 | 0.367-0.728 | 0.612 |
| IL-12B | 0.843 | 0.730-0.956 | 1.1E-4 | 0.553 | 0.367-0.738 | 0.575 |
| CCL19 | 0.699 | 0.521-0.878 | 0.024 |  |  |  |
| IL-10 | 0.810 | 0.683-0.938 | 4.5E-4 | 0.578 | 0.391-0.764 | 0.407 |
| TNFB | 0.797 | 0.667-0.927 | 0.001 | 0.590 | 0.408-0.772 | 0.336 |
| CXCL10 | 0.826 | 0.703-0.948 | 2.3E-4 | 0.748 | 0.586-0.909 | 0.008 |

Figure S1

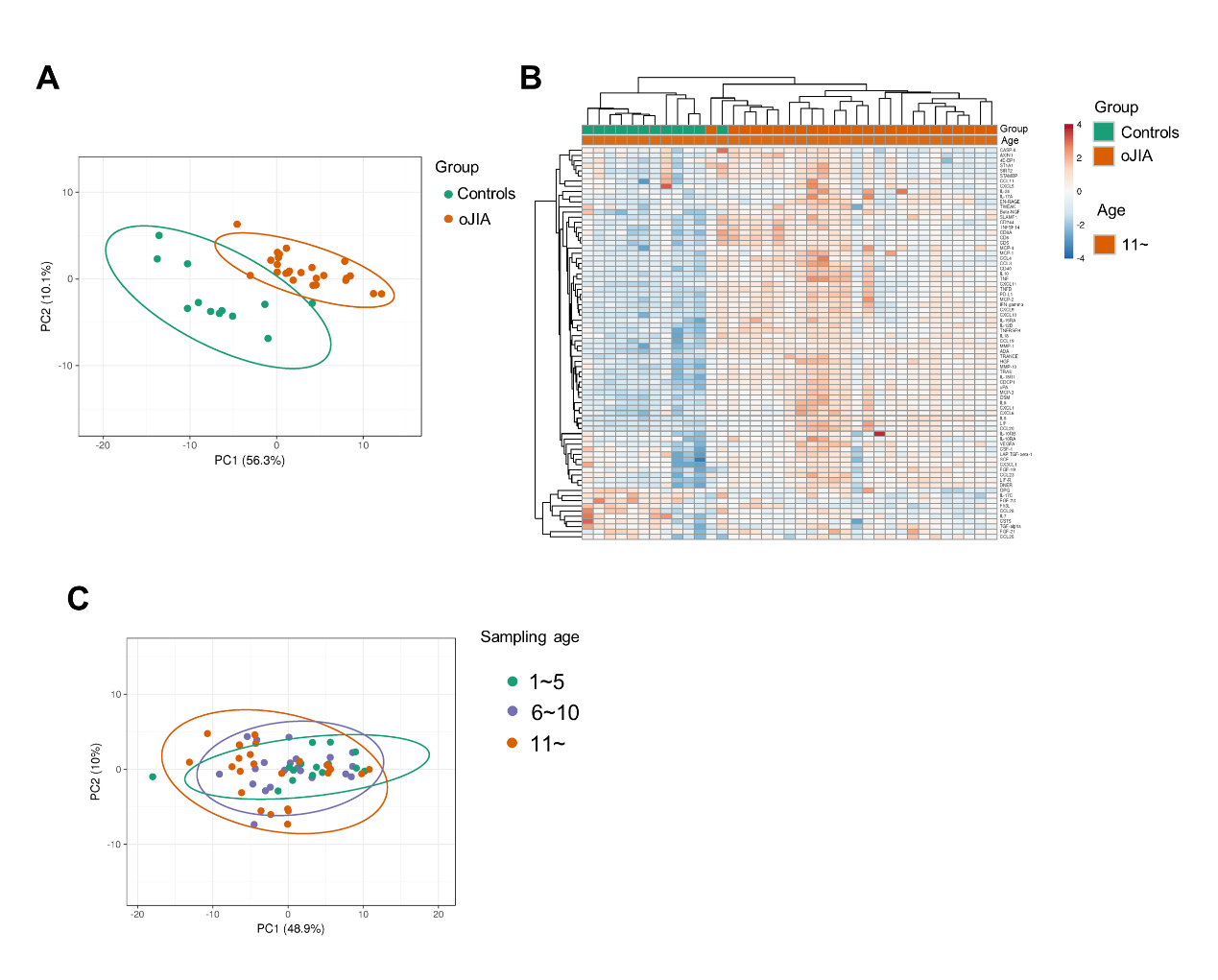

Fig S1. Analyses results of oJIA patients over 11 years old and controls. A) PCA results of the oJIA(>11y) and control groups based on 76 included proteins in SF present a separation between two groups. Confidence level of the ellipses is 0.95. Each point represents a single patient, with a total number is 37 including 25 oJIA and 12 controls. B) Heatmap visualization of 76 included proteins expression in oJIA and control group. C) PCA results of the oJIA with three different age group based on 76 included proteins in SF present an overlap among three group. Confidence level of the ellipses is 0.95.

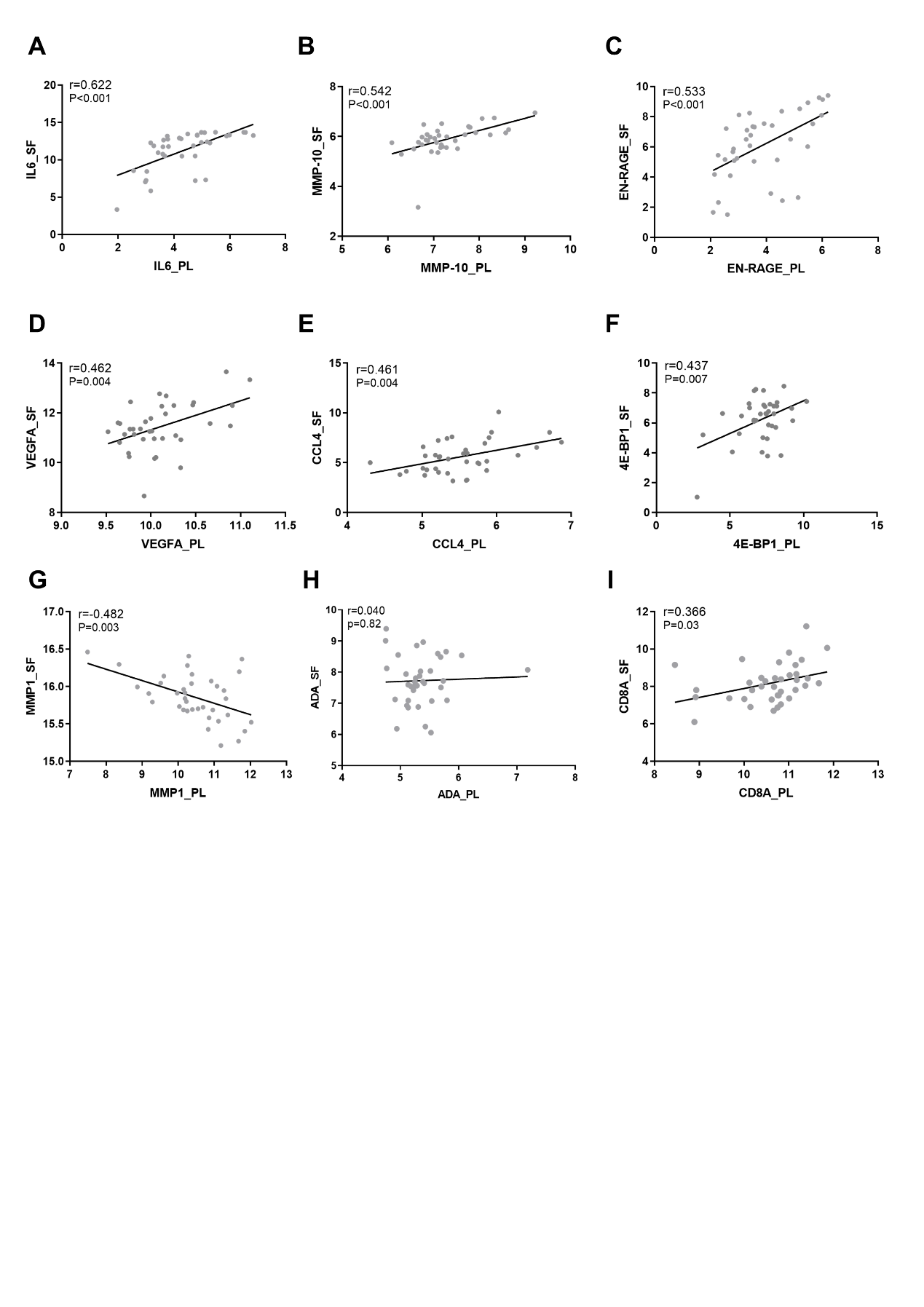

Figure S2. Correlations of DEPs between plasma and SF. A-G) Specific scatter plots of the proteins that exhibit significant correlations between plasma and SF (n=37). Pearson correlation analysis was applied to A-G. H) The scatter plot shows no correlation of ADA between plasma and SF. I) The scatter plot shows a weak correlation of CD8A between plasma and SF.

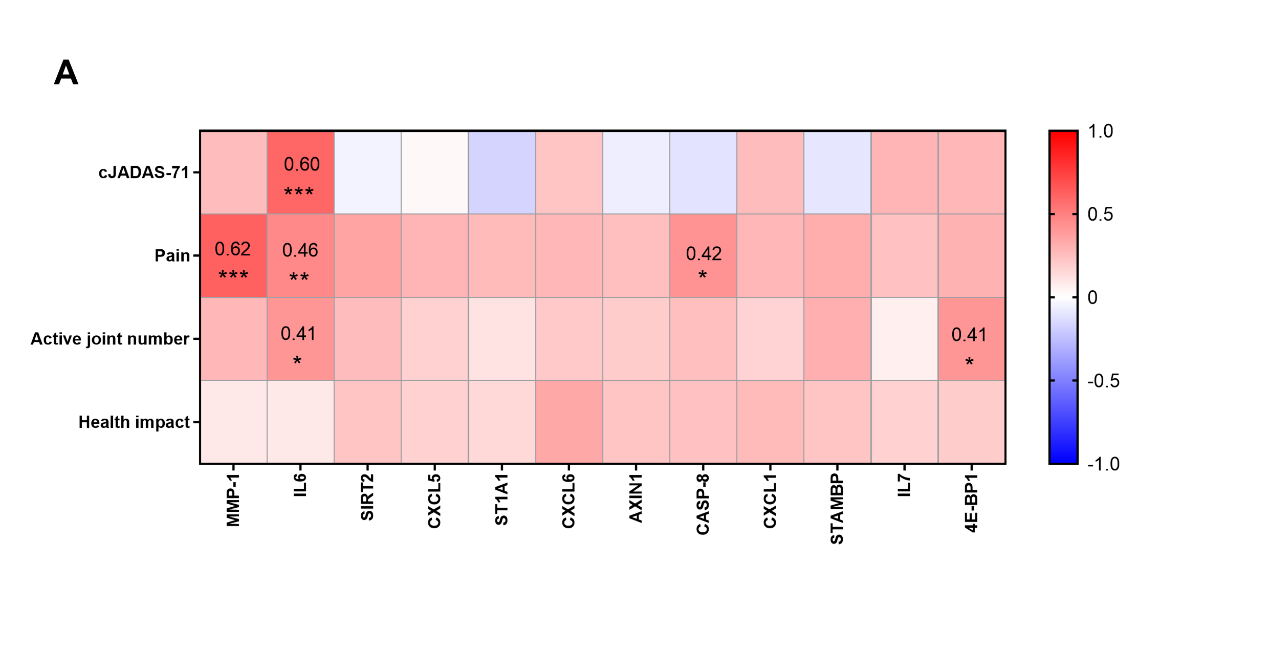

Fig S3. Correlations between DEPs in plasma and selected clinical parameters. A) Heatmap visualization of the correlations between DEPs in plasma and selected clinical parameters (cJADAS-71, pain, active joint number, and health impact). * P < 0.05, ** P < 0.01, *** P < 0.001
